# Emergence and clonal expansion of *Vibrio aestuarianus* lineages pathogenic for oysters in Europe

**DOI:** 10.1101/2022.04.04.486953

**Authors:** Aurélie Mesnil, Maude Jacquot, Céline Garcia, Delphine Tourbiez, Lydie Canier, Audrey Bidois, Lionel Dégremont, Deborah Cheslett, Michelle Geary, Alessia Vetri, Ana Roque, Dolors Furones, Alison Garden, Petya Orozova, Isabelle Arzul, Mathieu Sicard, Guillaume Charrière, Delphine Destoumieux-Garzón, Marie-Agnès Travers

## Abstract

*Crassostrea gigas* oysters represent a significant global food source, with 4.7 million tons harvested per year. In 2001, the bacterium *V. aesturianus francensis* emerged as a pathogen that causes adult oyster mortality in France and Ireland. Its impact on oyster aquaculture has increased in Europe since its reemergence in 2012. To better understand the evolutionary mechanisms leading to the emergence and persistence over time of this pathogen, we conducted a survey of mollusk diseases through national reference laboratories (NRLs) across Europe. We analyzed 54 new genomes of *V. aestuarianus* (*Va*) isolated from multiple environmental compartments since 2001, in areas with and without bivalve mortalities. We used a combination of comparative genomics and population genetics approaches to show that *Va francensis* lineages have undergone clonal expansion in Europe, likely after a recent selective bottleneck. Low mutation and recombination rates may have selected particular virulent genotypes. Furthermore, we identified a specific *cus-cop*-containing island conferring copper resistance to *Va francensis* whose acquisition may have favored the emergence of pathogenic lineages adapted to oysters.

## Introduction

Marine populations are under increasing threat of emerging infectious disease (EID) (1), contributing to mass mortality events impacting corals, sea-stars, mollusks and fish (2–4). Aquaculture is practiced in an open and highly connected environment and thus particularly threatened by pathogen propagation (5). Moreover, aquacultural practices frequently result in high population densities and many animal transfers, increasing the risk of pathogen introduction and dispersal (6).

Oysters (*Crassostrea gigas*) are frequently impacted by bacterial pathogens of the genus *Vibrio* (7–9). Vibrios are ubiquitously distributed in the oceans and show a wide range of niche specialization, ranging from free-living forms to those attachment to biotic and abiotic surfaces, from symbionts to pathogens, and from estuarine inhabitants to deep-sea piezophiles living under high pressure (10). Among them, *Vibrio aestuarianus* subsp. *francensis* (*Va francensis*) has caused recurrent mortality of adult oysters since 2001 (11). Having previously been restricted to France, this disease has now spread across European countries (France, Italy, Ireland, and Spain) (12, 13). This spread of *Va francensis* has resulted in mortality rates of approximately 25% and is a major economic concern as it impacts market-size oysters (14). Previous field studies found that pathogenic strains of *Va francensis* can colonize sentinel oysters, but are rare in samples from the local environment, *i.e.* sediment and water column. These data suggest that oysters are the preferred habitat of this pathogen (15). Moreover, it has been proposed that *Va francensis* can persist in oyster tissues during winter (15). These observations suggest *Va francensis* has adapted to oysters and raise questions about how this host specialization has occurred.

The emergence of pathogenic bacterial lineages can follow the acquisition of new traits which increase the ability to colonize a host (16). Trait acquisition can be achieved by mutation (17), but it occurs more frequently through horizontal transfer (through transformation, conjugation or transduction) of one or more genes favoring rapid adaptation to new niches and hosts (18). After emergence, the dynamics of pathogenic lineages are dictated by the relative importance of evolutionary forces: mutation, selection, genetic drift and recombination. Populations whose evolution is not or little influenced by recombination evolve clonally (19). One of the factors limiting the rates of horizontal transfer and recombination in a population is ecological isolation (20, 21) which limits opportunity to exchange genetic material with other populations.

Previous studies on *V. aestuarianus* diversity were based on a limited number of genomes, not reflecting overall diversity across diverse isolation contexts (22). This lack of genomic data hinders a complete understanding of the mechanisms governing the emergence and evolutionary dynamics of pathogenic lineages in *Va francensis*.

In the present study, we analyzed 54 genomes of isolates collected since 2001 across Europe. We combined phylogenetic and population genetic approaches to decipher evolutionary processes associated with the origin of the emergence of this oyster pathogen. Our study shows that Va *francensis* strains exhibit numerous features commonly reported in specialized pathogens. In particular, our data suggest the acquisition of genes involved in copper resistance is one mechanism promoting *V. aestuarianus* adaptation to oysters.

## Results

### European spread of *V. aestuarianus* oyster pathogens

To assess the geographic distribution of pathogenic *V. aestuarianus* in Europe, a questionnaire was sent in 2021 to all European laboratories in charge of the diagnosis and surveillance of mollusk diseases (supplementary material). Eight out of the 12 respondent laboratories reported that *V. aestuarianus* was detected during adult oyster mortality events. The earliest detections were reported by France and Ireland in 2001, and the most recent detection was reported by Denmark in 2021 (case used DNA detection only) (fig. 1, A). The number of countries reporting detection has increased since 2011, with 8 European countries currently reporting *V. aestuarianus*. Importantly the date of first detection may be under-estimated, since surveillance and testing for the pathogen has varied between countries and was only implemented in the majority of the laboratories following the development of a diagnostic tool in 2009 (23).

**Figure 1 :**
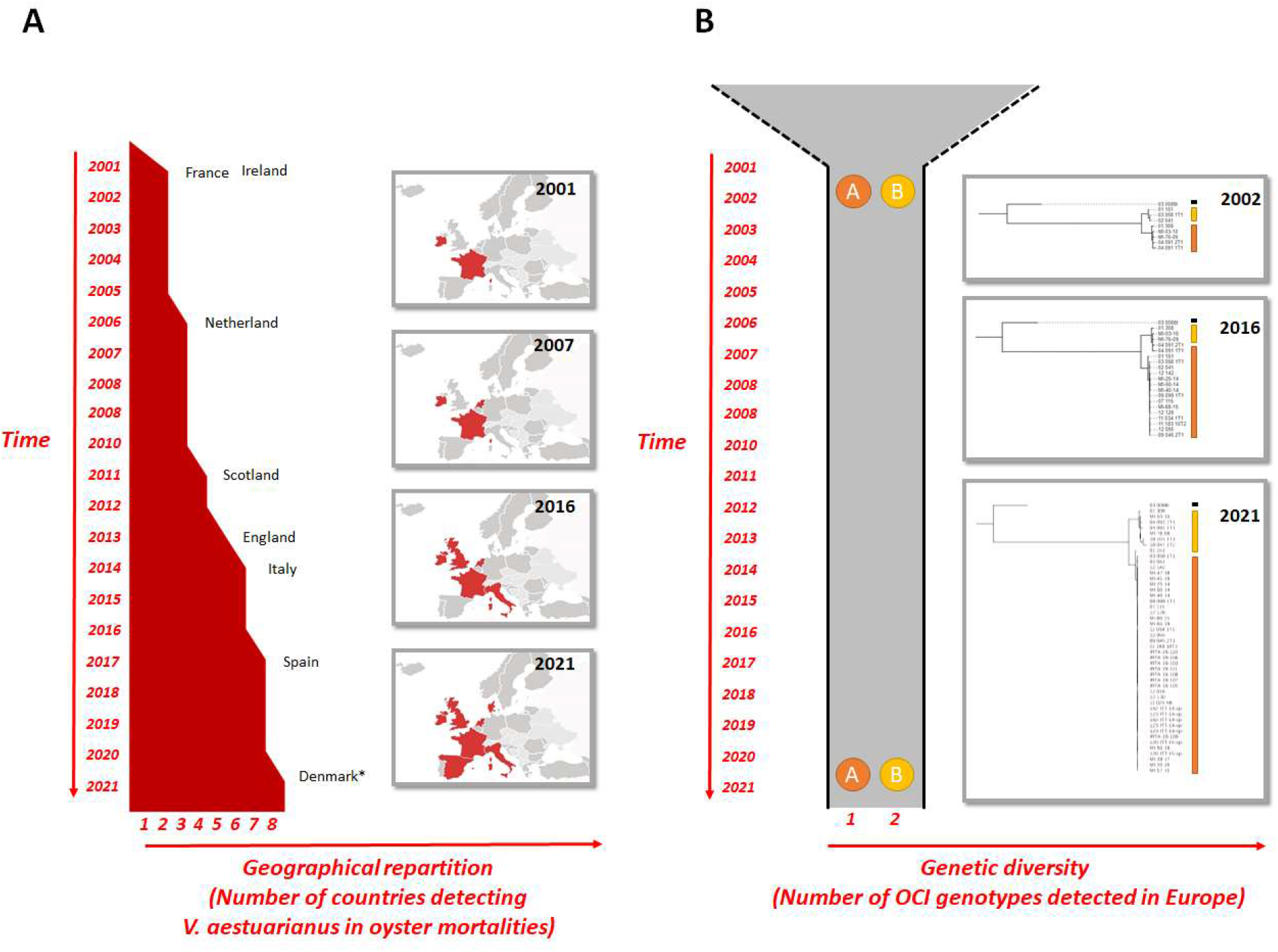
Figure 1: European expansion of V. aestuarianus francensis. While the geographic distribution of V. aestuarianus strains involved in C. gigas mortalities expands, genetic diversity remains stable over time. A) The number of countries reporting the detection of V. aestuarianus was assessed through a questionnaire sent to European national reference laboratories (NRLs). *Note: Denmark case report corresponds to a suspected case without bacterial isolation. B) The two lineage expansions: 2 genotypes were determined based on a maximum likelihood phylogenetic tree from complete genome alignments (GTR substitution model).

### *V. aestuarianus* diversity is structured according to environmental origin and isolation context

In order to assess the overall population structure of *V. aestuarianus* in Europe, we studied genetic diversity at the whole genome level using *V. aestuarianus* isolates from a broad variety of origins. 54 strains were sequenced and genomes were *de novo* assembled. We obtained 15 4.5-Mb complete genomes (with 2 to 4 contigs) and 39 draft genomes (with up to 150 contigs) (table S1). Whole genomes were aligned, and relationships between isolates were studied by building a maximum likelihood phylogeny (fig. 2). Overall, genomes were close to each other in the tree and showed highly similar nucleotide compositions, showing average nucleotide identities (ANIs) higher than 97.1% for all pairwise comparisons (table S2). The tree showed a tightly clustered clade of oyster clinical isolates (*i.* e., isolates from moribund oysters) (n = 25) inside genetically diverse *V. aestuarianus* strains isolated outside of oyster mortalities (n = 29), encompassing non-clinical isolates (i.e. strains isolated from sites without mortalities) and strains isolated during cockles mortalities. All oyster clinical isolates are encompassed in *Va francensis* subspecies (24) and all *Va francensis* isolates were found in oysters, and showed higher virulence levels when assayed through injection into oyster muscle (p-value< 0.001). The reconstructed tree also confirmed that this subspecies is composed of two lineages named A (n = 17) and B (n = 8) (22).

**Figure 2 :**
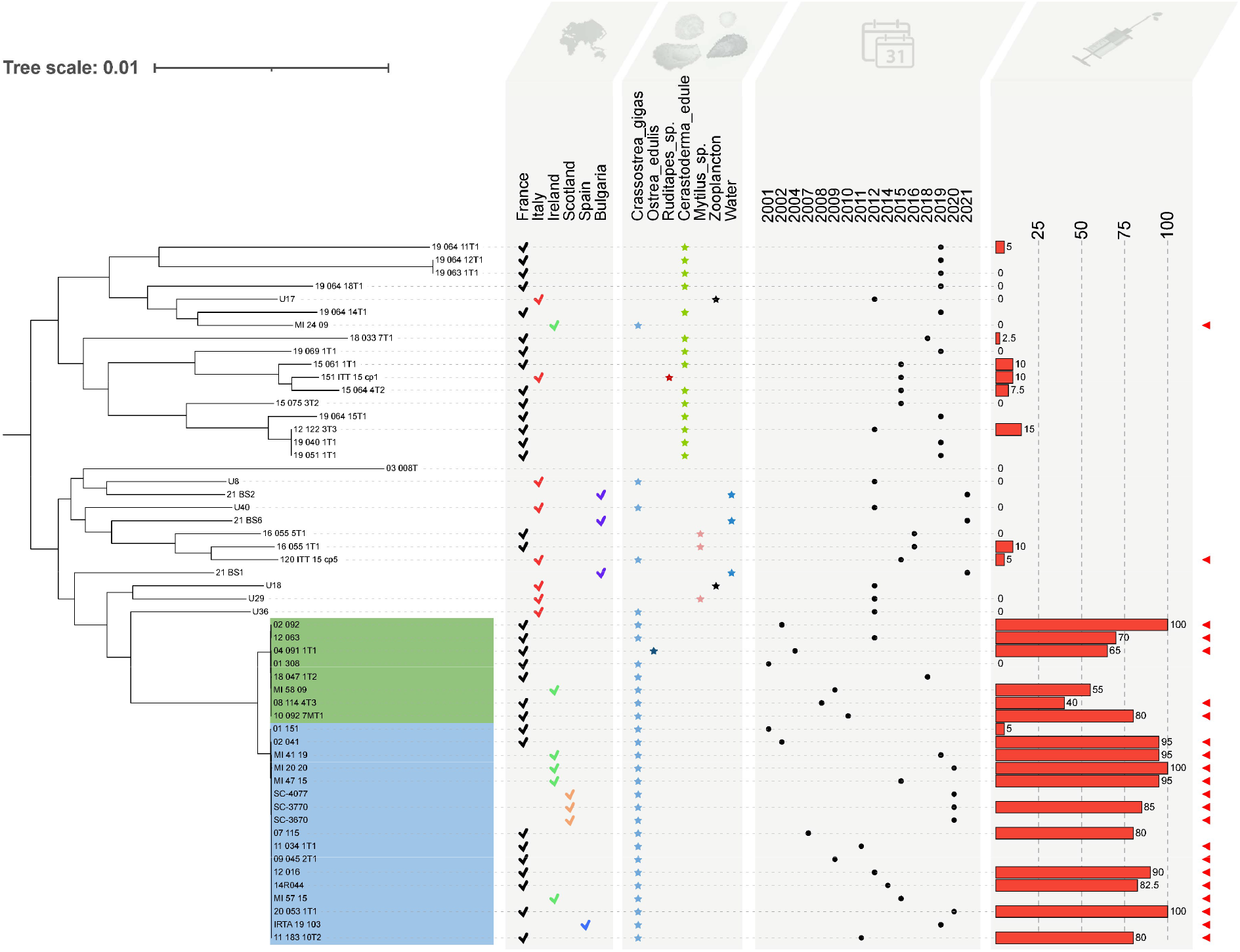
V. aestuarianus diversity is structured according to host species and isolation context. Phylogenetic tree of V. aestuarianus species calculated with RAxML using a GTR substitution model and rooted with the midpoint rooting option from whole-genome alignments. Several layers of annotations are added: country (check mark), matrix of isolation (star), year of isolation (point), virulence testing in the laboratory by injection into oysters (107 UFC/animal, duplicates of 10 animals) indicated with a bar plot on the right side representing mortality percentages, and context (strains isolated during oyster mortality events are indicated with a red arrow on the right). Va francensis lineages A and B are highlighted in blue and green, respectively. Their virulence is significantly higher compared to non clinical isolates (i. e. strains isolated on sites without mortalities) by Wilcoxon-Mann-Whitney test (p-value < 0.001). The absence of virulence data means the absence of a test for this strain.

Analysis of molecular variance (AMOVA) performed on the sequence dataset revealed that the isolation context (“oyster mortality” or not) and environmental origin (five host species, zooplankton and water) explained most significantly (p < 0.005) genetic variation (table 1). AMOVA also indicated the absence of spatial and temporal delineation of the oyster pathogen lineages. Altogether, our data reveal *V. aestuarianus* lineages linked European expansion of this oyster pathogen.

**Table 1 :** Analysis of molecular variance supporting genetic partitioning according to environmental origin and isolation context

| Factor | Variability (%) | p-value |
| --- | --- | --- |
| **Environmental origin** | **17.8** | **0.002** |
| **Isolation context** | **16.0** | **0.001** |
| Country of isolation | 7.1 | 0.074 |
| Year of isolation | 3.4 | 0.25 |

### Temporal emergence of oyster pathogen lineages

Next, we used a Bayesian approach to date the emergence and reconstruct the temporal evolution of oyster pathogens lineages. As a critical prerequisite, we checked whether the *Va francensis* subspecies is a measurably evolving population. A temporal signal was observed with a correlation between genetic distances from a reconstructed common ancestor and strains sampling time, yielding a R^2^ of 0.4236 (fig. S1A). The best-fitting evolutionary model for the *Va francensis* population was obtained under a GMRF Bayesian skyline population model with a relaxed clock, leading to a rate of 7.27 × 10^-7^ substitutions per position per year, or 3.2 mutations per genome per year. The mean time of emergence of the most recent common ancestor (TMRCA) of the lineage A was estimated to be 1956 (95% highest posterior density (HPD): 1919–1972). At this time, a divergence occurred between strain 01-151, which is non-virulent toward oysters, and other lineage A strains that are mostly pathogenic. The TMRCA of lineage B maps to the late 1970s, albeit with uncertainty (95% HPD: 1898–1993). According to the coalescence-based reconstruction of demographic variation, the *Va francensis* population has slowly decline since its emergence, with accelerated decline since 2000 (fig S1B).

### Genome stability in *V. aestuarianus francensis* lineages

Recombination can drive genetic diversity and evolutionary dynamics in bacterial populations. We therefore quantified homologous recombination events using *V. aestuarianus* genome alignments.

#### Recombination has little impact on oyster pathogen evolution

We found that recombination (r) has a lower impact on the core genome substitution rate in *Va francensis* lineages than the impact of mutation (m), with mean r/m = 0.36+/− 0.22 and 0.28+/− 0.09 in lineages A and B respectively. The ratios of these lineages are significantly (student test p-value < 0.005) smaller than in non-clinical *V. aestuarianus* isolates (i.e. strains isolated from sites without mortalities) (r/m = 0.91 +/− 0.45). 7.8 and 3.3 recombination events were identified on average for these lineages. Recombination in genomes can generate phylogenetic inconsistencies between taxa that can emerge in phylogenetic networks as the presence of alternative pathways (25). The network that we constructed from the whole-genome alignment (fig. S2) confirmed that recombination does not affect *Va francensis* isolate relationships, with few ambiguities among these taxa. One known consequence of the absence of recombination impact is strong linkage between loci in non-recombinant genomes (26). In *Va francensis* lineage A and, to a lesser extent, in lineage B, strong linkage between single nucleotide polymorphisms (*SNPs*) was observed (fig. S3). In contrast, the recombination impact detected in all other genomes was confirmed with phylogenetic networks, which showed ambiguities between taxa (fig. S2) and low linkage between loci in these genomes (fig. S3). However, recombination might have supported the emergence of the *Va francensis* group: 488 recombination events were detected at the node corresponding to the hypothetical common ancestor of all pathogenic oyster isolates (fig. S4).

#### Core genome stability was observed in *Va francensis* lineages

Mutations also generated little genetic diversity among *Va francensis*. Indeed, apart from the recombinant segments, among the 19 775 variant genomic positions in the full dataset, 666 and 83 positions vary among *Va francensis* A and B lineages. Finally, low impact of recombination and mutation on *Va francensis* lineages results in core genome stability. We used Tajima’s D to compare observed genetic diversity in *V. aestuarianus* to that expected under neutral evolution and constant population size. Deviation from neutrality was larger among the *Va francensis* lineages A (Tajima’s D = −2.3) and B (−2.5) than in the full dataset (−0.7), with less polymorphism than expected under neutral evolution (fig. 3 A). To determine whether the low genetic diversity was due to demographic effects (affecting whole genomes) or to selection effects (affecting particular genes), we adopted a gene-based approach to investigate selection pressures at the gene level. While most of the 1,974 core genes were under neutral evolution outside of the *Va francensis* subspecies, the vast majority were monomorphic within lineages A and B (fig. 3 B), which is consistent with the purging of genetic diversity through bottlenecks during transmission dynamics.

**Figure 3 :**
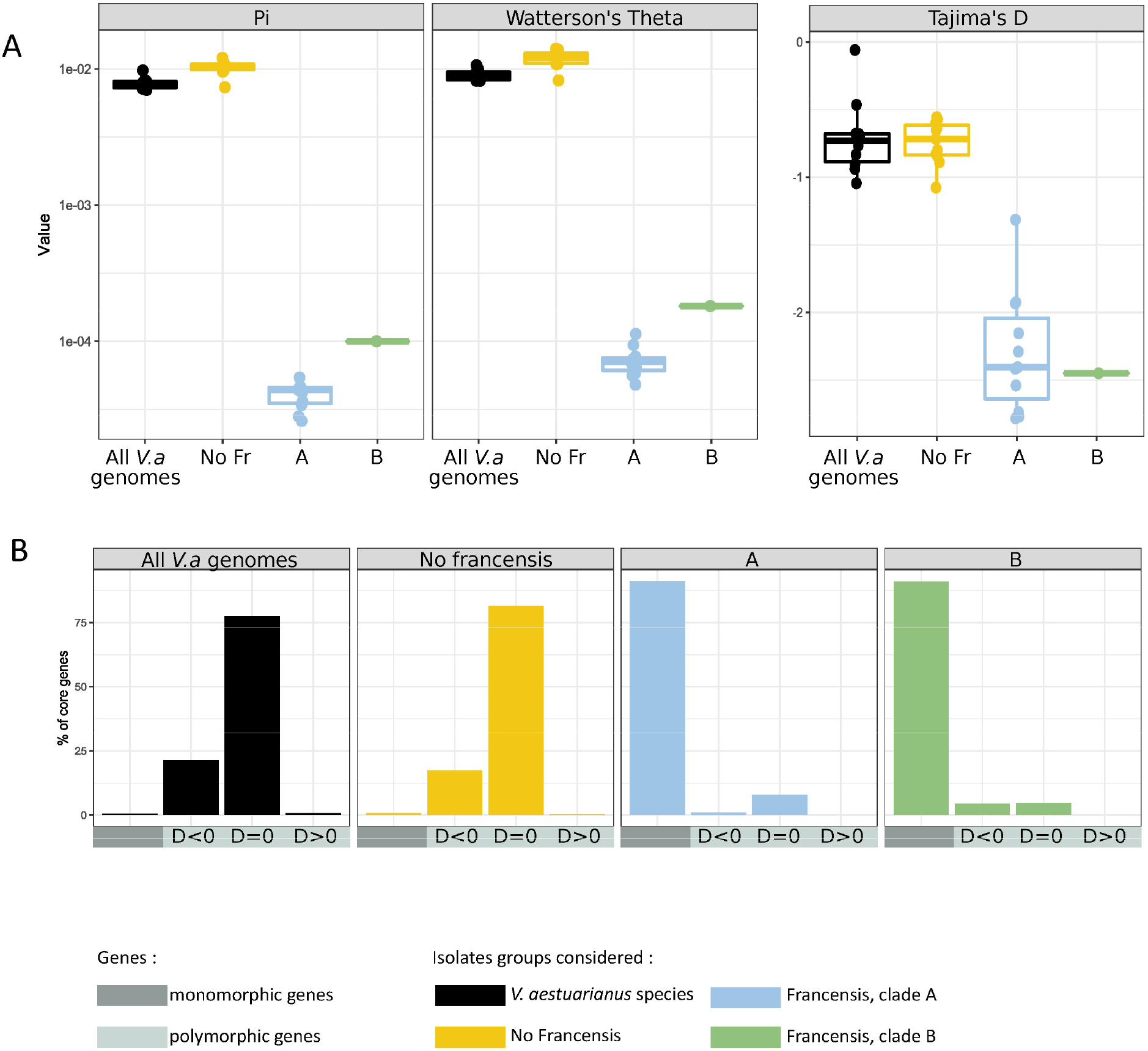
Support for a recent bottleneck that purged genetic diversity in V. aestuarianus francensis lineages. Genetic statistics in V. aestuarianus species (black), other than Va francensis (yellow), Va francensis, lineage A (blue) and lineage B (green). A) Whole genome scale. Even through the expected genetic diversity under neutral evolution and a stable population size in Va francensis is low (Watterson’s theta), the observed diversity (Pi) is still lower, as shown by the Tajima’s D value below - 2. Pi: observed genetic diversity; Theta_w: Watterson estimate of expected genetic diversity; Tajima_D: Tajima’s D test statistic. The Y axis in the Pi and Watterson’s theta plots is log10 scaled. B) Gene scale. Core genes show different selection patterns among groups of strains and among core genes within groups. While most core genes are under neutral evolution in isolates other than Va francensis, most genes are monomorphic in Va francensis lineage A and B. D>0 and D<0: significantly negative or positive Tajima’s D value (p < 0.05) and D=0 (p > 0.05). A negative Tajima’s D is indicative of purifying selection or recent population expansion, while a positive Tajima’s D is indicative of diversifying selection or population contraction. A zero value is indicative of neutral evolution.

#### *Va francensis* subspecies were found to have a closed pangenome

with genes largely shared among all isolates and almost no strain-specific genes, as determined by dataset pangenome analysis (fig S5). In contrast, other *V. aestuarianus* isolates exhibited many strain-specific genes. Altogether, our data support the clonal evolution of the two *Va francensis* lineages.

### Positive selection signatures of oyster pathogen lineages

We used *pcadapt*, which detects structure and clustering of individuals in a population, to detect potential signatures of positive selection affecting oyster pathogen lineages. 2,925 SNPs were selected with an adjusted p-value <0.001. Based on principal component analysis, principal component 1 (PC1) segregated the *Va francensis* lineages from the rest of *V. aestuarianus* strain collection (fig. S6). 46 outlier SNPs were associated with PC1 including 17 non-synonymous and one stop-gained SNP potentially involved in diversifying selection. These genes are listed in table S3 and notably include genes encoding flagellar proteins (*pomA*) or related to chemiotaxis.

### Oyster-pathogenic *V. aestuarianus* have genomic features of specialist pathogens

According to low rates of homologous recombination and the closed pangenome, the two *Va francensis* lineages can be considered as clonally evolving lineages. Furthermore, clonal evolution, as well as the small number of known habitats (15), suggest that these lineages are oyster specialists. Pathogen populations have often undergone a change of niche (passage from a free-living lifestyle to an obligatory parasitic one), ultimately resulting in ecological isolation and thus inhibition of DNA transfer with the parental population, or even with the broader bacterial community (27, 28).

We further tested the hypothesis of specialization by analyzing genomic features known to be enriched during transition to specialist lifestyle : insertion sequences (ISs) and pseudogene (29). We found a higher number of ISs in the genomes of pathogenic oyster lineages A and B than in strains isolated from contexts without bivalve mortality (p-value = 0.01 and 0.02, respectively). Interestingly, in groups containing strains isolated during cockle mortality events, ISs were more abundant than in pathogenic oyster genomes (p-value = 0.02 and 0.03, respectively) (fig. 4). IS families were conserved between oyster pathogen genomes, with most ISs belonging to the IS66, IS902 and ISAS1 families (table S4 and fig S7). IS conservation in theses clades suggests that IS enrichment and expansion in genomes stopped after pathogen emergence. The dynamics of the ISs appeared to be different in the cockle isolates. Indeed, the number and families of ISs represented appeared to be poorly conserved. We suspect that IS flux is active in cockle isolates, with ongoing horizontal exchange and intragenomic expansion.

**Figure 4 :**
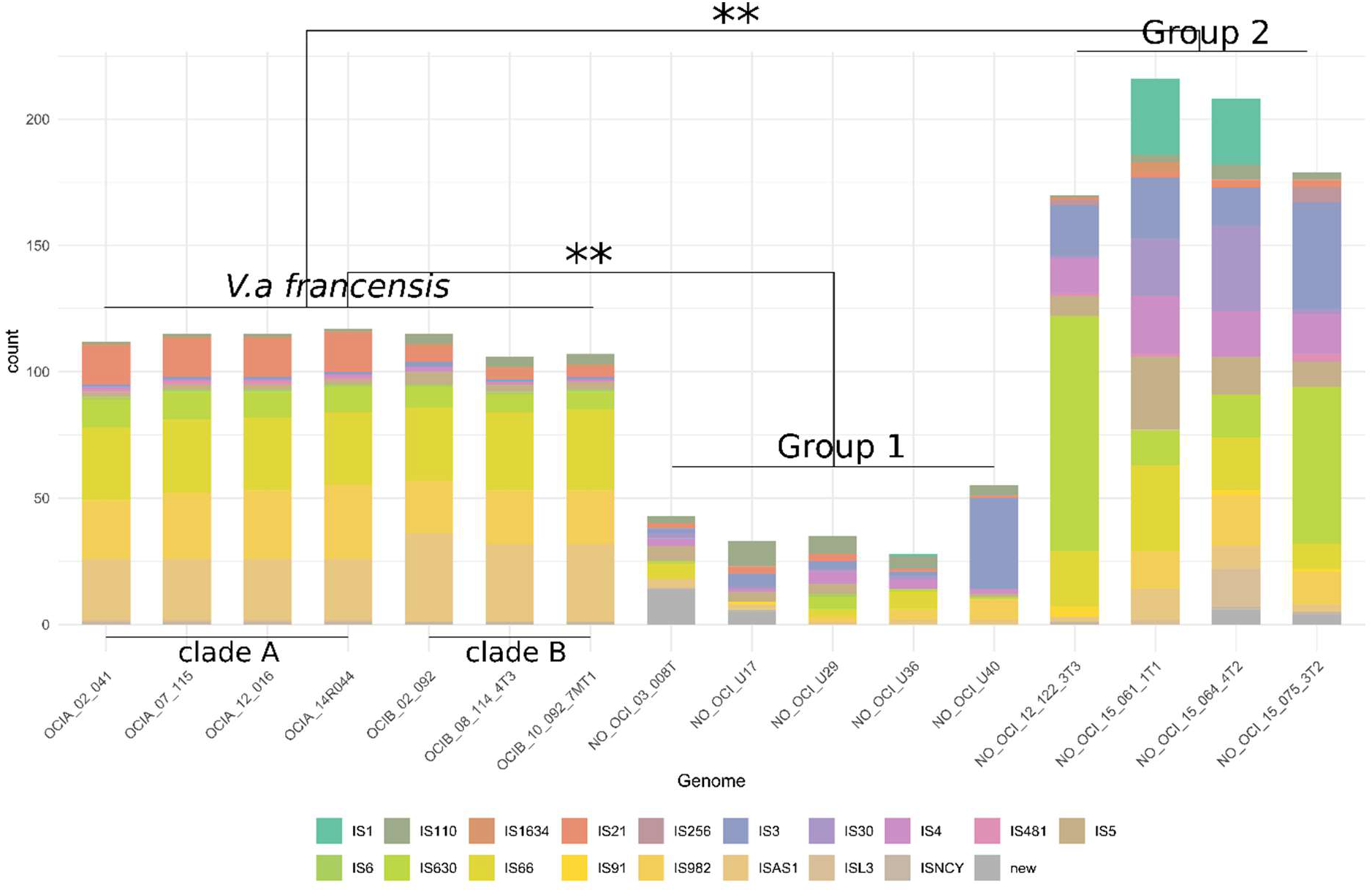
Differential Insertion sequence (IS) enrichment is observed in Va francensis and cockle associated isolates genomes. Distribution of the number of IS elements (complete and partial) in 16 complete genomes (chromosomes A and B). Colors represent IS families predicted by ISEScan. The genomes are those of 4 Va francensis lineage A strains, 3 Va francensis lineage B strains, 5 non clinical strains (isolated on sites without mortalities) (group 1), and 4 strains isolated during cockle mortality events (group 2). The conservation of IS number and families between genomes of Va francensis from lineage A or clade B indicates an ancestral origin of ISs. The genomes that contain the most ISs are those of strains isolated during cockle mortality events (group 2). The diversity of IS quantities and families represented suggests that IS enrichment is in progress. The number of ISs in groups was significantly different between Va francensis and other strains from group 1 and group 2 (**: Kruskal test p value < 0.05).

To check whether IS enrichment is involved in pseudogenization, we crossed IS and pseudogene genomic coordinates and determined the number of ISs with pseudogenes around them (from 2,000 bp before the IS start to 2,000 bp after the IS end) (table S5 and S6). Regardless of the genome and chromosome considered, the vast majority (on average 86%) of ISs had pseudogenes nearby. IS and pseudogene enrichment supports our hypothesis that the *Va francensis* lineages are specialized toward oyster.

### Copper resistance acquisition: One potential step in the emergence of oyster-pathogenic clades

Trait acquisition is often the first step preceding selective bottlenecks and clonal expansion of host-adapted pathogens. To identify traits that may have preceded oyster pathogen emergence, we studied the *V. aestuarianus* pangenome and detected 139 genes that were shared exclusively by *Va francensis* genomes. Most of them (106) were annotated as hypothetical proteins. The remaining 33 genes encode proteins with functions related to copper export, cell-bacteria interaction (LPS/antigen O, cell wall or peptidoglycan), chemotaxis and DNA repair. The sequences of these 139 genes were identical or well conserved between strains (table S7).

Some oyster pathogen specific genes localize to a putative mobile genetic element (MGE), as determined by crossing their coordinates with IS coordinates in an oyster pathogen genome (02_041) (fig. 5 A). This putative MGE contains Cus operon genes (*cusABCF,S/R*) surrounded by ISs from the IS66 family (length = 15 kb), as well as the copper export genes *copA* and *copG* and the regulator *cueR*, located near tn7 transposon genes (tnA, B, C and E) (length = 10 kb), and is later referred to as a *cus-cop*-containing island.

**Figure 5 :**
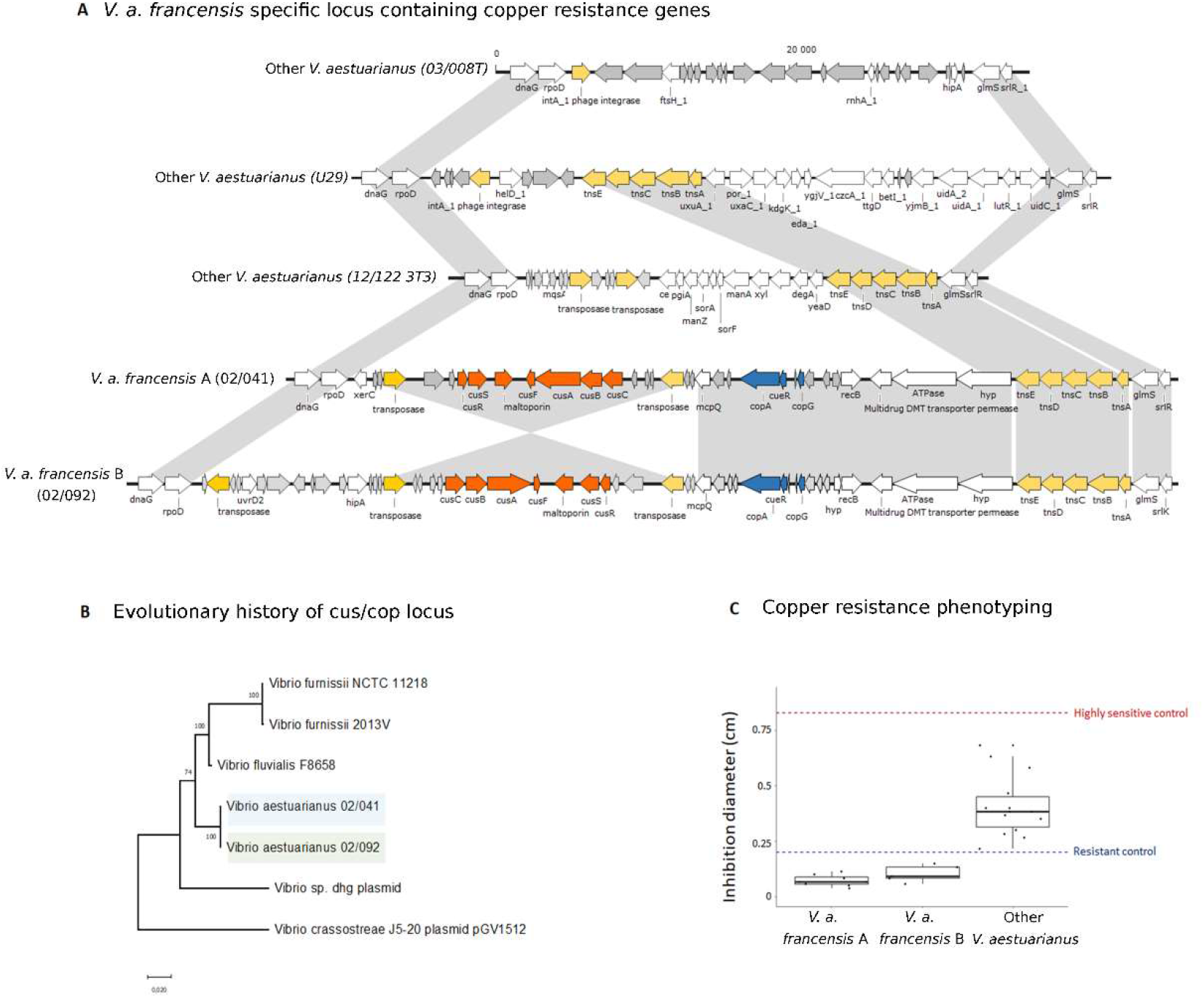
A) Va francensis-specific locus containing copper resistance genes. Transposase and integrase are indicated in yellow, cus genes in orange, and cop genes in blue. B) Evolutionary history of the cus/cop locus. NJ, bootstrap test (500 replicates). The evolutionary distances were computed using the Kimura 2-parameter method and are shown as the number of base substitutions per site. This analysis involved 8 nucleotide sequences. All ambiguous positions were removed for each sequence pair (pairwise deletion option). There was a total of 29 555 positions in the final dataset. Evolutionary analyses were conducted in MEGA X [4]. C) Agar plate assay: bacterial growth inhibition diameter around a pellet impregnated with 50 mM CuSO_4_ solution. Va francensis linages A and B are significantly more resistant to copper that other V. aestuarianus strains (Wilcoxon-Mann-Whitney test p value < 0.001).

The *Va francensis* ancestor could have acquired copper export genes by horizontal transfer. Presence in an MGE could have facilitated integration of these genes into the bacterial genome. To investigate the origin of *cus-cop*-containing islands, we used BLAST analyses of each operon and of the entire region in 127 reference genomes of different *Vibrio* species (table S8 and fig S8). The *cus-cop*-containing island presence is scattered along the *Vibrio* phylogeny and is not specific to a monophyletic group. It appeared highly similar to some *V. fluvialis* and *V. furnissii* genomic sequences with a nucleotide identity >95%. Moreover, this region was close to plasmidic elements identified in *V. crassostreae* and *Vibrio* sp. dgh with a nucleotide identity >89% (fig. 5B).

To test whether the genomic differences between the oyster pathogens and other strains could lead to phenotypic differences, we tested copper resistance in 27 *V. aestuarianus* strains using an agar diffusion assay. Supporting our hypothesis, all tested oyster pathogens (8 strains from lineage A and 5 from lineage B) appeared significantly (p-value < 0.001) more resistant than other strains to copper in radial diffusion assays (fig. 5C).

## Discussion

Here, we describe a genomic analysis of *V. aestuarianus* isolated from multiple European countries, in different epidemic contexts and from multiple environments (five host species, zooplankton and water) since 2001. We report the expansion and clonal evolution of two oyster-pathogenic lineages from the *V. aestuarianus francensis* subspecies (11).

We estimate that 7.23 x 10^-7^ substitutions occur per site per year in *Va francensis*, similar to the evolutionary rate inferred in *V. cholerae* during pandemic waves (30): this indicates that pathogenic *Va francensis* lineages emerged during the 1990s, a decade when global oyster production greatly increased according to Food and Agriculture Organisation. Indeed, oyster production increased to over 140,000 tons per year in the 1990s (31). According to the reconstructed demographic history, the effective population size of *Va francensis* is small and showed a slowly decrease that accelerated in the 1990s. This sharper decreased may have followed emergence of oyster-pathogenic lineages, indicating a possible bottleneck that resulted in selection of oyster-adapted strains. This bottleneck may have purged polymorphisms from this ancestral population, explaining the observed monomorphism in *Va francensis* core genes. Indeed, some pathogenic monomorphic lineages may endure population bottlenecks, supporting acquisition of new functions and colonization of new hosts (32).

Several genomic modifications may help pathogenic oyster lineages adapt to their host, including core genome substitutions and gene acquisition. We found non-synonymous substitutions that were fixed in pathogenic oyster lineages. These occurred in genes involved in host interaction or virulence, such as the *pomA* gene which encodes a flagellum motor protein and the gene *luxP* linked to quorum-sensing. The functional consequences of these substitutions remain to be explored.

Acquisition of copper resistance traits may also have contributed to the emergence of the oyster-pathogenic lineages. Our data indicate that the *Va francensis* ancestor acquired a genomic island carrying copper detoxification genes. Indeed, copper is a heavy metal found at high levels in coastal waters because of agricultural and industrial activities (33–35). Copper can accumulate in oyster tissues (up to 18,000 μg/g in China (36)), particularly at infection sites (37) where it supports antimicrobial defense as copper can kill invading microorganisms (37–39). Bacteria have, in turn, developed copper detoxification mechanisms which utilize the *cop* and *cus* genes (40). *Cop* genes support initial Cu^+^ translocation from the cytoplasm to the periplasm*. Cus* genes encode a CusABC system that exports periplasmic copper into the extracellular medium (41). Importantly, *copA* is essential for copper resistance, oyster colonization and virulence in *V. tasmaniensis* LGP32, an oyster-pathogenic strain associated with the Pacific Oyster Mortality Syndrome (42). Similarly, *copA* deletion in *V. crassostreae* also attenuates virulence in oysters (7). Our data further support copper resistance as a shared trait that allows pathogens to adapt to oysters as a host.

Our results suggest clonal evolution in *Va francensis* lineages. Recombination has a low impact on genetic diversity. The r/m ratio, ~0.3 in both lineages, is similar to clonally evolving pathogenic bacteria like the *V. cholerae* 7 pandemic lineage (r/m = 0.1 ; (43)), *M. tuberculosis* complex (between 0,426 and 0,565, (44)) and lineage 1 of *Salmonella enterica* (r/m = 0.2 ; (45)). We also found that *Va francensis* has a closed pangenome. This led us to formulate the hypothesis of an ecological isolation of these lineages, a hallmark of specialized pathogens that do not exchange genes with other bacterial populations. This hypothesis is also supported by enrichment of insertion sequences and pseudogenization in both oyster-pathogenic lineages. During specialization, pathogens transition from a free-living lifestyle to a permanent association with a host. Through host association, pathogens can acquire compounds of intermediate metabolism that allow genes associated with related biosynthetic pathways to become facultative or redundant (25). Studies on pathogen specialization have largely focused on highly specialized intracellular bacteria, which can undergo an extreme genome rationalization process (26). This process implies pseudogenization of non-essential genes that accumulate mutations or ISs and can ultimately result in gene loss (29, 46, 47). However, genome size reduction was not observed, probably because *Va francensis* pathogenic lineages are not obligate intracellular parasites. While oyster pathogens have a restricted habitat, they can still replicate in the absence of their host, at least in rich medium. Moreover, *Va francensis* can colonize diverse host tissues, including hemolymph, gills and digestive connective tissues (15). As a consequence, *Va francensis* is adapted to diverse tissular micro-environments, consistent with being less specialized than obligate intracellular bacteria. Further studies of oyster pathogen microhabitats may help to elucidate their complete intra-host life cycle, and the functions of specific genes linked to adaptation. Whether clades A and B occupy similar niches in infected oysters and whether they have the potential to cooperate remains unknown. Neither epidemiological parameter estimation (48) nor in-depth analysis of their genomes identified lineage-specific functional traits.

Here we report the clonal expansion of *Va francensis* lineages across Europe. It has been reported that another major pathogen, the herpes virus Ostreid OsHV-1, has also caused the mortality of *C. gigas* (49) along the French coasts since the emergence of a new variant in 2008 (50). Double infections by OsHV-1 and *Va francensis* were reported in French oyster farms in 2014 and 2015 (51, 52) and have also been observed in Ireland since 2008 (Cheslett, pers comm). Although experimental infections combining the two pathogens at high doses in spat and juvenile *C. gigas* have not demonstrated direct collaboration between them (53, 54), at the population level the virus may still weaken animal defenses and thus amplify the impact of *Va francensis*.

The global spread of pathogens is a recurrent problem in agriculture and aquaculture and is often favored by anthropogenic introductions via infected hosts (55). The many transfers taking place during the lifetime of an oyster probably explain most of the spread of pathogenic lineages of *Va francensis* across Europe. Preventing the spread of pathogens during transfers is particularly complicated in the case of *Va francensis* since the animals may be asymptomatic (15). There is therefore an urgent need to develop sensitive detection assays to monitor the spread of pathogens: the specific *Va francensis* genes revealed in this study, such as the cusS/R genomic island, may provide new diagnostic tools, which may help to monitor and prevent future mortality events.

## Material & methods

### Isolate selection

The study collection was designed to represent the temporal, geographic and clinical heterogeneity of *V. aestuarianus* origins. However, ascertainment bias could occur due to sampling strategy used. Indeed, strains were mainly isolated during bivalve mortalities. We maximized recovery of the genetic diversity of oyster pathogenic *V. aestuarianus* by collecting isolates from all European countries reporting mortalities related to *V. aestuarianus* or presence of the bacteria. We also included strains isolated during field surveys of sites without known mortality events. Strains were collected with the help of European NRLs responsible for the diagnosis and surveillance of mollusk diseases: Ifremer (France), MI (Ireland), IZSVE (Italy), IRTA (Spain), Marine Scotland Science (Scotland), and NRLFMC (Bulgaria). Sequence comparisons of isolates from France, notably those isolated from moribund oysters, turned out to be very similar. In order to equilibrate our sampling, we excluded 17 isolates identical with included isolates. We selected approximately one strain per batch of moribund oysters (a batch represents a sampling point) (table S9). The 54 isolates used in the study are summarized in table S10.

### Illumina, Ion torrent and PacBio whole-genome sequencing

Genome sequencing was performed between 2019 and 2021. Several sequencing technologies were used depending on the objectives. We used a long-read sequencing technology for some strains to improve assembly quality and study genomic features. DNA was isolated from 35 strains using the NucleoSpin Tissue Kit (Macherey-Nagel) following the manufacturer’s guidelines. The concentration of each sample was adjusted to 0.2 ng/μl for library preparation using the Illumina Nextera XT DNA Library Preparation Kit according to the manufacturer’s guidelines. For each library, paired-end reads of 150 bp were produced on the Illumina NextSeq 550 platform (Bio Environnement, Perpignan, France). The quality of the resultant fastq files was assessed using FastQC (56). Primers, Illumina adapter sequences, low-quality bases at the 3’ end of reads with Phred-scaled quality scores less than 30 and reads with a length <50 bp were removed using Trim Galore (57). Quality-controlled reads were *de novo* assembled using the SPAdes genome assembler v3.13.1 (58) with the default kmer size and a coverage cutoff value of 20 reads (59).

For 2 strains (04_091_1T1 and U8), enzymatic shearing was performed on 500ng of genomic bacterial DNA using the NEBNext Fast DNA Library Prep kit (New Englands Biolabs) and following the manufacturer’s recommendations. Samples were barcoded with individual IonCodes (Thermo). Purified libraries were then quantified by real-time PCR using the Ion Universal Library Quantification Kit (Thermo) prior to be diluted and pooled at 25μM each. The Ion Torrent S5 libraries were prepared using the “Ion 510™ & Ion 520™ & Ion 530™” for Ion Chef Kit and sequenced on the Ion Torrent S5 using an Ion 530 semi-conductor sequencing chip (Thermo).

DNA was isolated from other strains using a MagAttract HMW DNA Kit (Qiagen) following the manufacturer’s instructions. Library preparation and sequencing on a PacBio Sequel II instrument were performed with the GENTYANE INRAe platform (Clermont-Ferrand, France). *De novo* genome assemblies were obtained using FLYE v2.8.3 (60). One round of polishing was performed using RACON v 1.4.3 (61).

The assembly was evaluated with QUAST, and genome completeness was checked using the BUSCO v5.1.1 (62) and vibrionales_odb10 databases as lineage datasets, including 57 Vibrionales as detailed in table S11.

Assembly features and GenBank accession numbers are summarized in table S1.

### Genome annotations

For each assembly, coding sequences (CDSs) were predicted using PROKKA v1.12 (63) with default databases and –compliant and –rfam flags and annotation of rRNAs, tRNAs and other ncRNAs enabled. Functional annotation of the detected CDSs was performed using EggNog-Mapper v2.1.0 (64, 65). ISs were detected in the genomes using ISEScan v1.7 (66). Pseudogene detection was performed with the GenBank files released from PROKKA with PSEUDOFINDER v1.0 (67), using the diamond flag and NCBI nonredundant database. These later analyses were performed only on resolved genomes to avoid obtaining false negatives in IS detection and false positives in pseudogene detection at the intercontig locations.

### Phylogeny and recombination analysis

Pairwise ANIs (%) were calculated using FastANI (68). Whole-genome alignments of the 54 isolate contigs were made using Progressive Mauve v13/02/2015 (69). Whole-genome alignment was used to infer the maximum likelihood phylogeny using RAxML v8.2.12 with the TVM model of nucleotide substitution (transversion model, AG=CT and unequal base frequencies) and 100 bootstrap replicates, with strain 03_008T as the outgroup.

To identify polymorphic sites, the haploid variant calling pipeline snippy v4.6 (available at https://github.com/tseemann/snippy) was used with default parameters.

To assess the influence of various structuring factors on the observed genetic diversity, we performed AMOVA on the sequence dataset, with environmental origin, isolation context, sampling year, and country as structuring factors, using the R package *pegas*.

Recombination was evaluated across this genome alignment using ClonalFrameML v1.12 (70). We used an extension that allows different recombination parameters to be inferred on different branches of the clonal genealogy. Linear regression analysis of the root-to-tip distances against sampling time was performed using TempEst v1.5.3 (71). The substitution rate was estimated under different demographic and clock models, using Beast v1.10.4, taking advantage of the sampling timeframe between 2001 and 2021. We tested both a strict and a relaxed molecular clock. Constant-sized, exponential-sized and GMRF Bayesian skyride plot models (72, 73) were used. For each model, two independent chains were conducted for 200 million generations and convergence was assessed by checking effective sample size values for key parameters using Tracer v1.7.1 (74). For robust model selection, both path sampling and stepping-stone sampling approaches were applied to each BEAST analysis to estimate the marginal likelihood (75, 76). The best model selected by Bayes factor comparison was a relaxed molecular clock with a GMRF Bayesian skyride demographic model. For this model, triplicate runs were combined using LogCombiner v1.10.4, with removal of 10% burn-in.

### Selection analysis

Tajima’s D was calculated considering core genome alignment using MEGA 11 Tajima’s test on neutrality. Core genome alignment were previously partitioned according to lineages or subspecies, resulting in *Va francensis* lineage A alignment (n = 17 sequences), *Va francensis* lineage B alignment (n = 8 sequences) and no *Va francensis* alignment (n = 29 sequences). Since sample size is known to influence population genetic parameter estimations (77), we calculated Tajima’s D on subsamples of eight sequences. For *Va francensis* lineage A and no *Va francensis* isolates, Tajima’s D was calculated on ten different alignments constructed with eight sequences randomly subsampled. Then we calculated mean of the Tajima’s D obtained from the ten alignments. For the *Va francensis* lineage B alignment, the eight sequences represented the entire sampled set. Tajima’s D was also calculated for each core genes, considering the same partitions, employing the R package *pegas*.

In order to detect SNPs that may explain *Va francensis* emergence, we performed a genome scan analysis using *pcadapt* package v.4.3.3 for R (78) that detect structure and clustering of individuals in a population. Number of principal components was set to six. We used LD.clumping option, with window size of 5 kb. LD.clumping uses p-value to sort the SNPs by importance (e.g. keeping the most significant ones). It takes the first one (e.g. most significant SNP) and removes SNPs that are too correlated with this one in a window around it. We also removed SNP with minor allele frequency under 0.05. After computing the test statistic, we used Bonferroni adjustment to correct for multiple comparisons, and threshold of 0.001 was applied to identify outliers SNPs.

### Bacterial virulence estimation by injection

Bacteria were grown with constant agitation at 22°C for 24 h in Zobell medium. Cultures were washed by centrifugation (10 min, 1500 g) and resuspended in artificial, sterile seawater to obtain OD_600nm_ = 0.1. Fifty microliters of bacterial suspensions (10^7^ cfu, checked by plating on Zobell agar) were injected intramuscularly into anesthetized specific pathogen-free (SPF) oysters produced and maintained in the IFREMER facility (79). After injection, the oysters were transferred to aquaria (10 oysters per 2.5 L aquarium, duplicate aquaria per bacterial strain) containing 2 L of aerated, UV-treated seawater at 22°C and kept under static conditions for 5 days.

### Copper resistance phenotyping

Paper discs (6 mm) were loaded with CuSO_4_ by soaking them overnight in 50 mM CuSO_4_. Cells from stationary phase cultures at DO = 1 were washed twice with SSW before being spread onto agar plates containing 15 g.L^-1^ bactopeptone and 0.5 M NaCl. Using sterile tweezers, copper-impregnated disks were placed on the bacterial lawn before incubation at 20°C. After 5 days of incubation, the diameter of the inhibition zone around the disk was measured.

### Statistic tests

Basic statistical analysis of all of the data were performed using R v4.1.2. Data were checked for normality and homogeneity of variance before analysis. Wilcoxon-Mann-Whitney was performed to evaluate if recombination ratios, IS number in genomes, virulence toward oysters and halo size (representing strains sensitivity to copper) differed among *Va francensis* lineages and between *Va francensis* lineages and other strains.

## Acknowledgments

We first thank the NRLs for mollusk diseases that contributed to the questionnaire survey (Bojan Adžić, Raquel Aranguren, Giuseppe Arcangeli, Deborah Cheslett, Marc Engelsma, Mark Fordyce, Ana Grade, Michael Gubbins, Lone Madsen, Petya Orozova, and Ewa Paździor). We thank Nicolas Bierne for helpful discussion and Viviane Boulo, Jean-Michel Escoubas, Nicole Faury and Gaelle Courtay for their valuable technical assistance. We also thank Bruno Petton and staff of IFREMER Bouin, who contributed to animal production and delivery. We thank the Bio-Environment platform (University of Perpignan Via Domitia) and Jean-François Allienne for support in library preparation and sequencing. We are grateful to the Ifremer Bioinformatics Core Facility (SeBiMER; https://ifremer-bioinformatics.github.io/) for providing technical assistance and scientific support in bioinformatics analysis. We also acknowledge the Pôle de Calcul et de Données Marines (https://wwz.ifremer.fr/en/Research-Technology/Research-Infrastructures/Digital-infrastructures/Computation-Centre) for providing DATARMOR computing and storage resources. This work was funded by Europe DG-Sante (through European Union Reference Laboratory for mollusk diseases funding) and French Ministry DPMA (MIRAGE project) and DGAl through NRL funding. Moreover, A.M., PhD, was supported by IFREMER and Région Nouvelle Aquitaine. This study is part of the « Laboratoire d’Excellence (LabEx) » TULIP (ANR-10-LABX-41) framework.

