## Supplementary figures for "Emergence and clonal expansion of *Vibrio aestuarianus* lineages pathogenic for oysters in Europe"

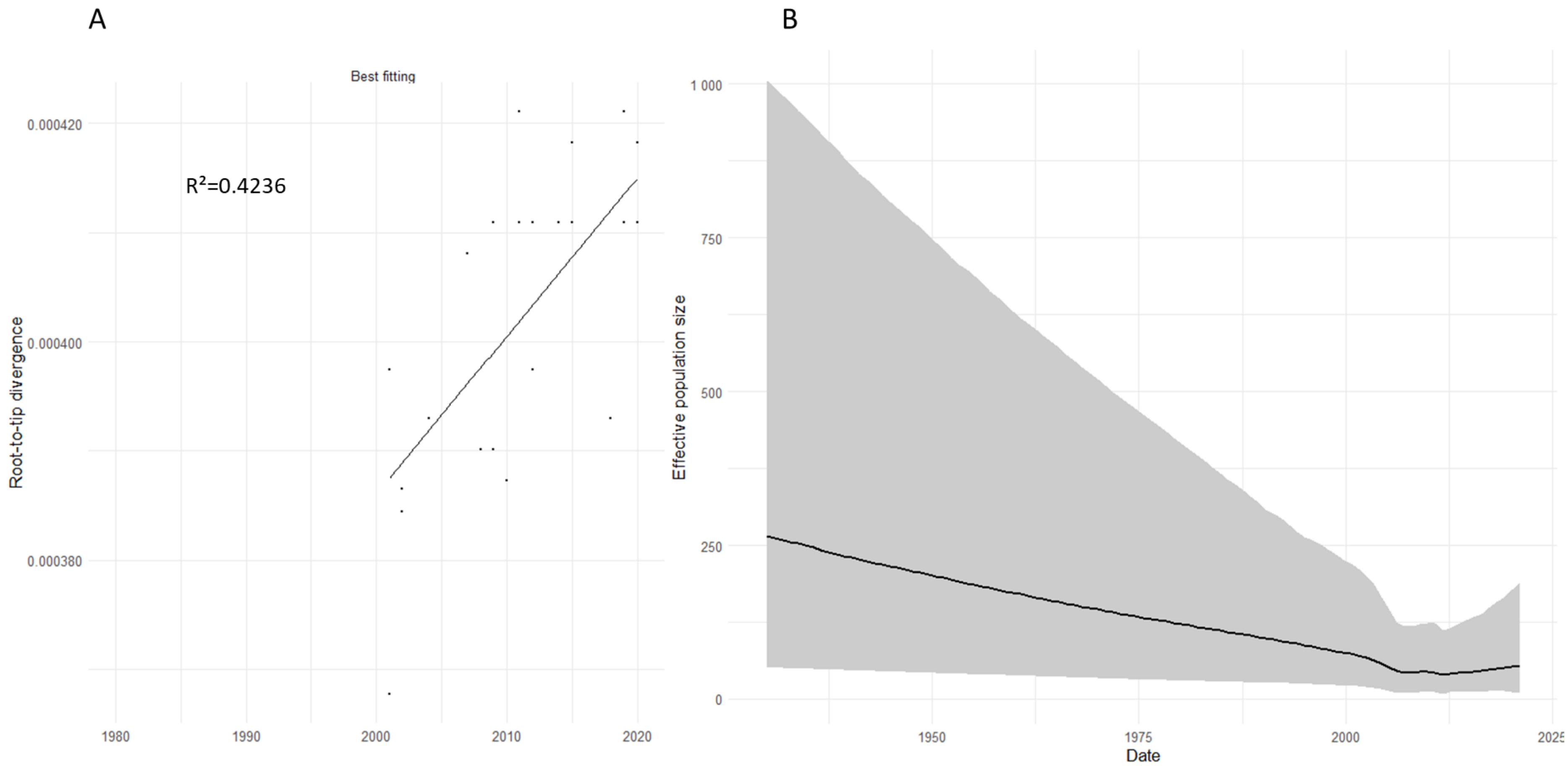


Figure S1: Temporal signal and reconstructed demographic history of *V. aestuarianus francensis*. A) Hierarchical root-to-tip analyses performed on the *V. aestuarianus francensis* data set, using TempEst (2007). B) GMRF Bayesian skyride plot indicating variation over time of the median of the effective population size of *V. aestuarianus francensis*. The estimated variations and the 95% confidence intervals are represented by black line and grey area respectively.

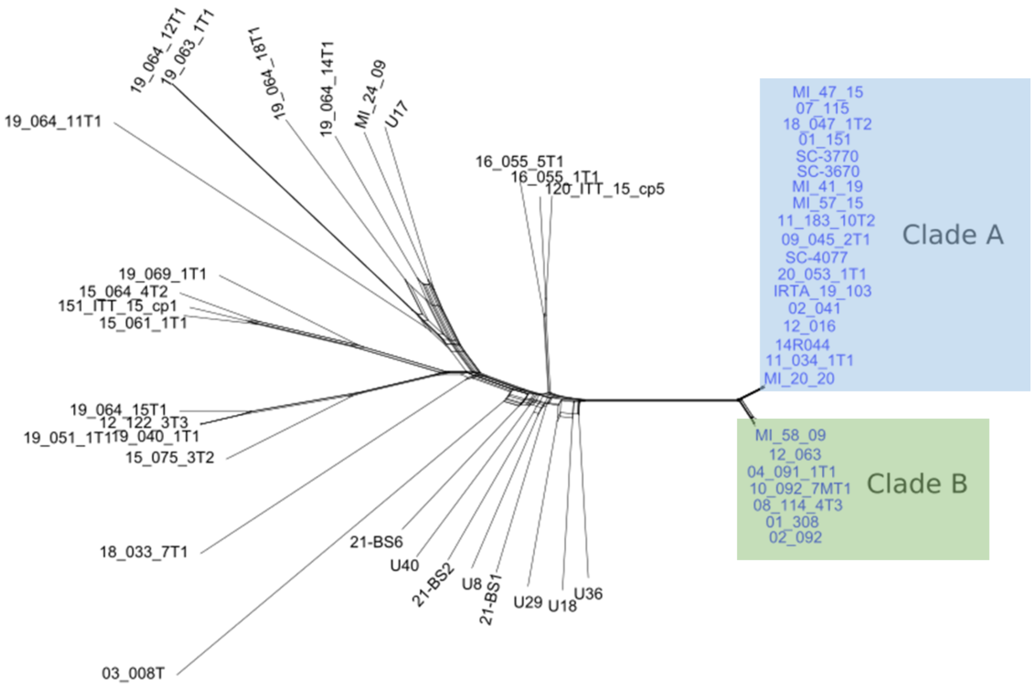


Figure S2: NeighborNet graph estimated from distance matrix calculated from the complete alignment. *Va francensis* lineages A and B are indicated in blue and green respectively

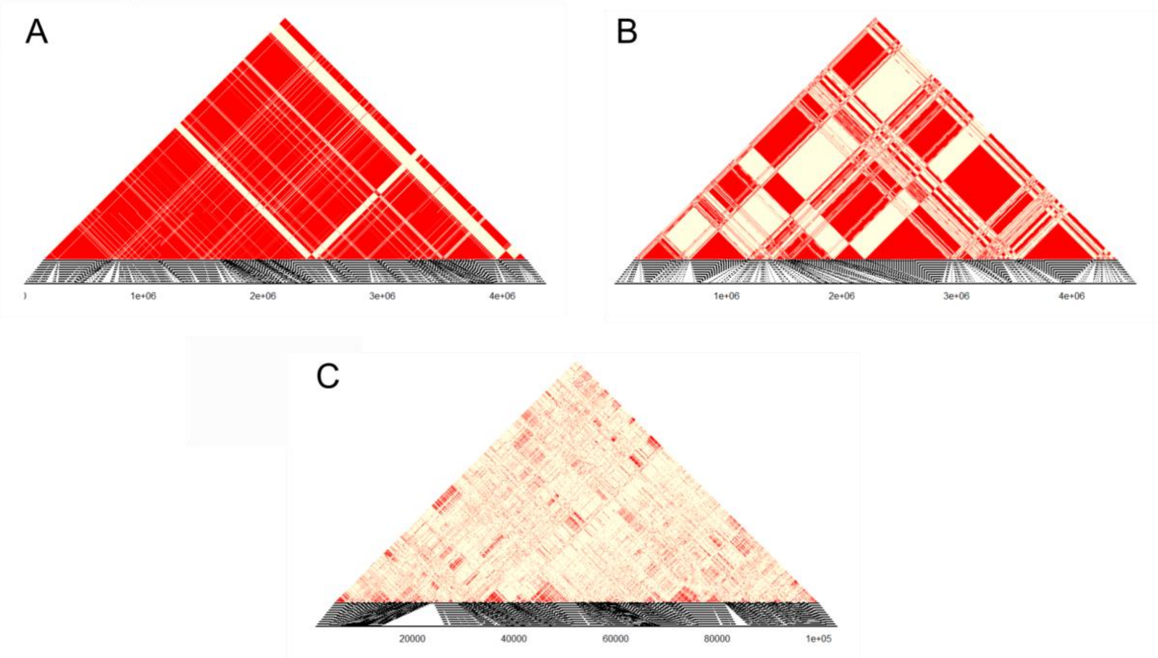


Figure S3: Association map between SNP in A) *Va francensis*, lineage A (n=17, all SNP); B) *Va francensis*, lineage B (n = 8, all SNP) and C) No francensis strains (n = 29, 5000 first SNP), obtained thanks to the LDscan and LDmap functions of the package *pegas* (R). The horizontal axis indicates the position of the loci on the chromosome. The linkage coefficients ($r^2$) between each pair of loci are indicated as coloured squares: the squares at the bottom of the triangle are for nearby loci, whereas the square at the top is for the two most distant loci. The stronger the association between two loci (*i.e.* found in all the sequences), the redder the square.

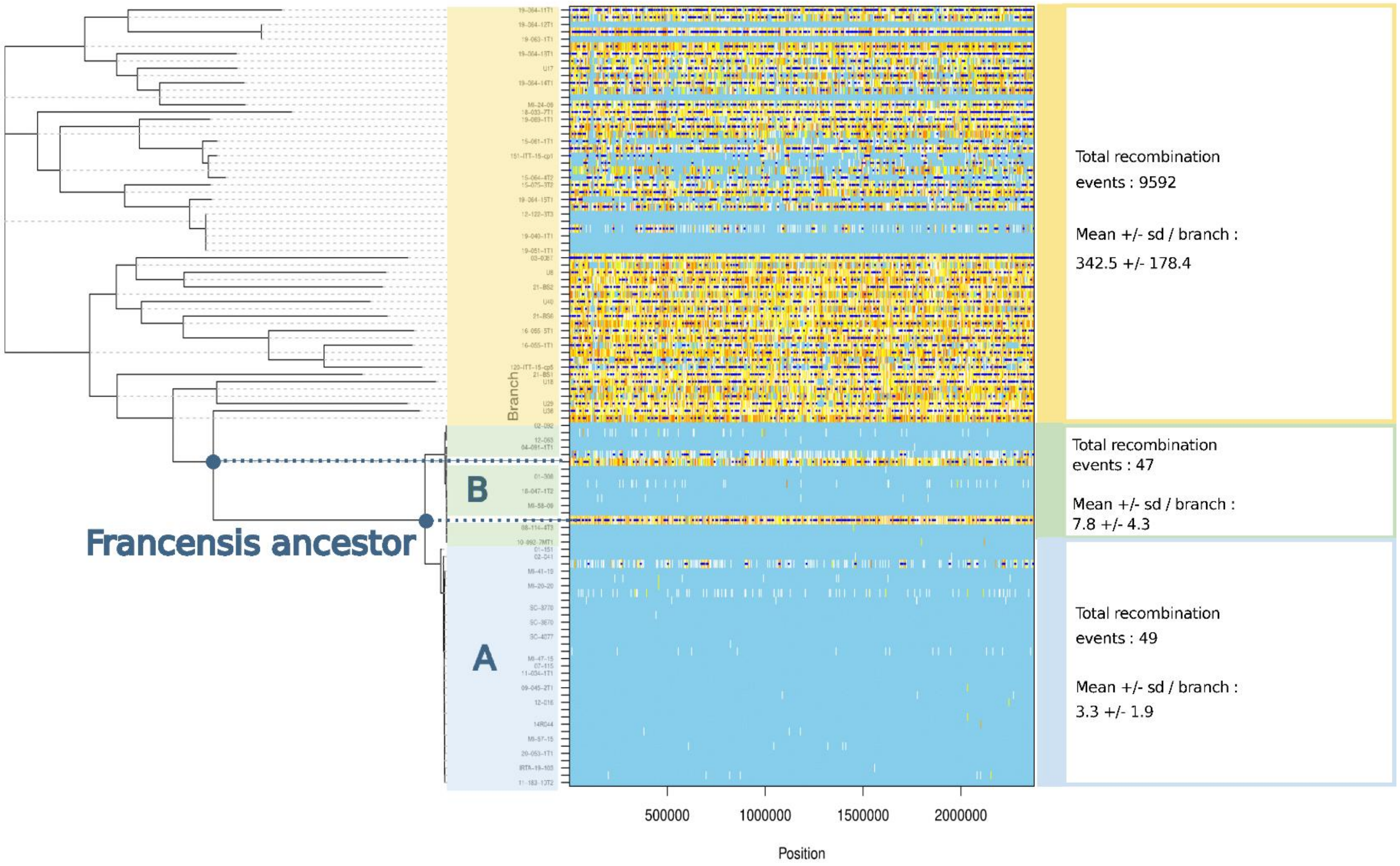


Figure S4: A limited number of recombination events occurred in *Va francensis* genomes after their expansion. Detected recombination events are shown by dark blue horizontal bars. For a given branch, light blue sites indicate no substitution. All other colors (from white to red) indicate a substitution. White represents a nonhomoplasic substitution. An increasing level of redness means an increasing degree of homoplasy. *Va francensis* lineages A and B are represented in blue and green, respectively, and the other strains are shown in yellow. For improved readability, some nodes of the tree as well as the row of the recombination events matrix related to these nodes are indicated with points and dashed lines.

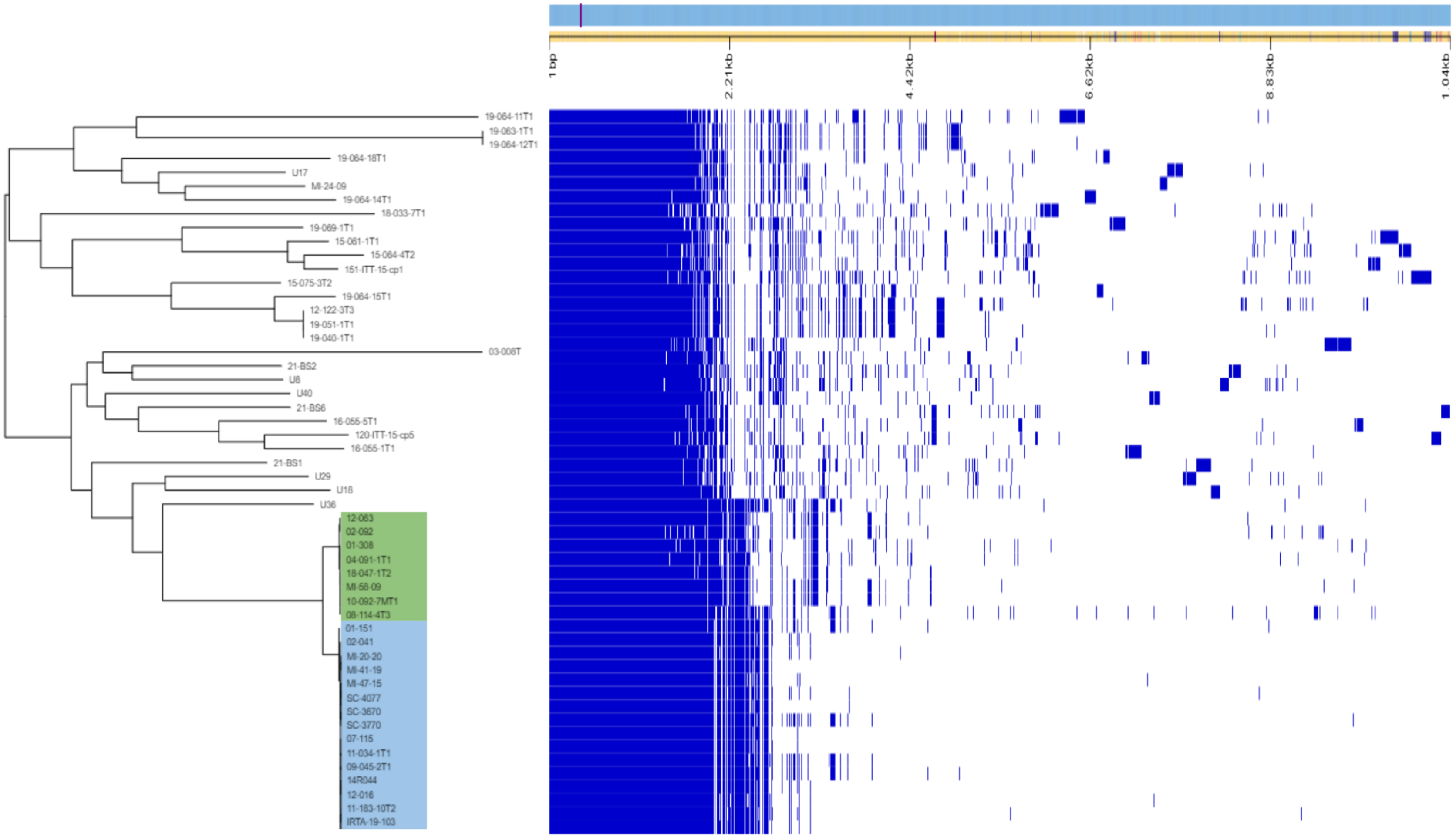


Figure S5: Phandango plot of Roary gene presence and absence. Genes are shown as light blue bricks along the top and are sorted left to right by the proportion of isolates they are observed in and might not represent the genome order. Presence (blue) and absence (white) of genes is plotted with respect to each isolate phylogenetic placement. *Va francensis* lineages A and B are represented respectively in blue and green color areas.

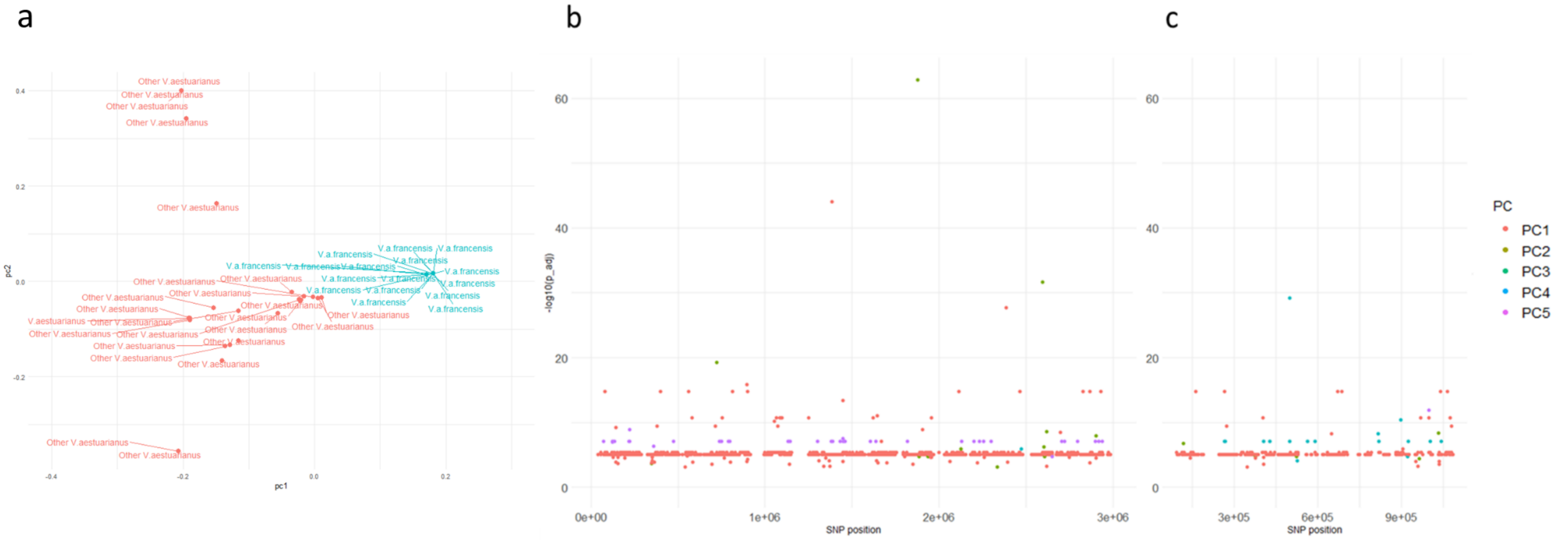


Figure S6: Genome scan analysis of *V. aestuarianus* strains for detecting SNPs involved in adaptation. A) Plot of the first two principal components (PC). The 54 *Vibrio aestuarianus* strains are represented by points and colorized according they belong to *Va francensis* (blue) or not (pink). B) and C) Manhattan plot respectively for chromosome 1 and 2, representing the 2925 SNPs and –log10(p.value). The SNPs are colorized according to the principal component to which they correlate most (PC1: pink, PC2: brown, PC3: green, PC4: blue, PC5: purple).

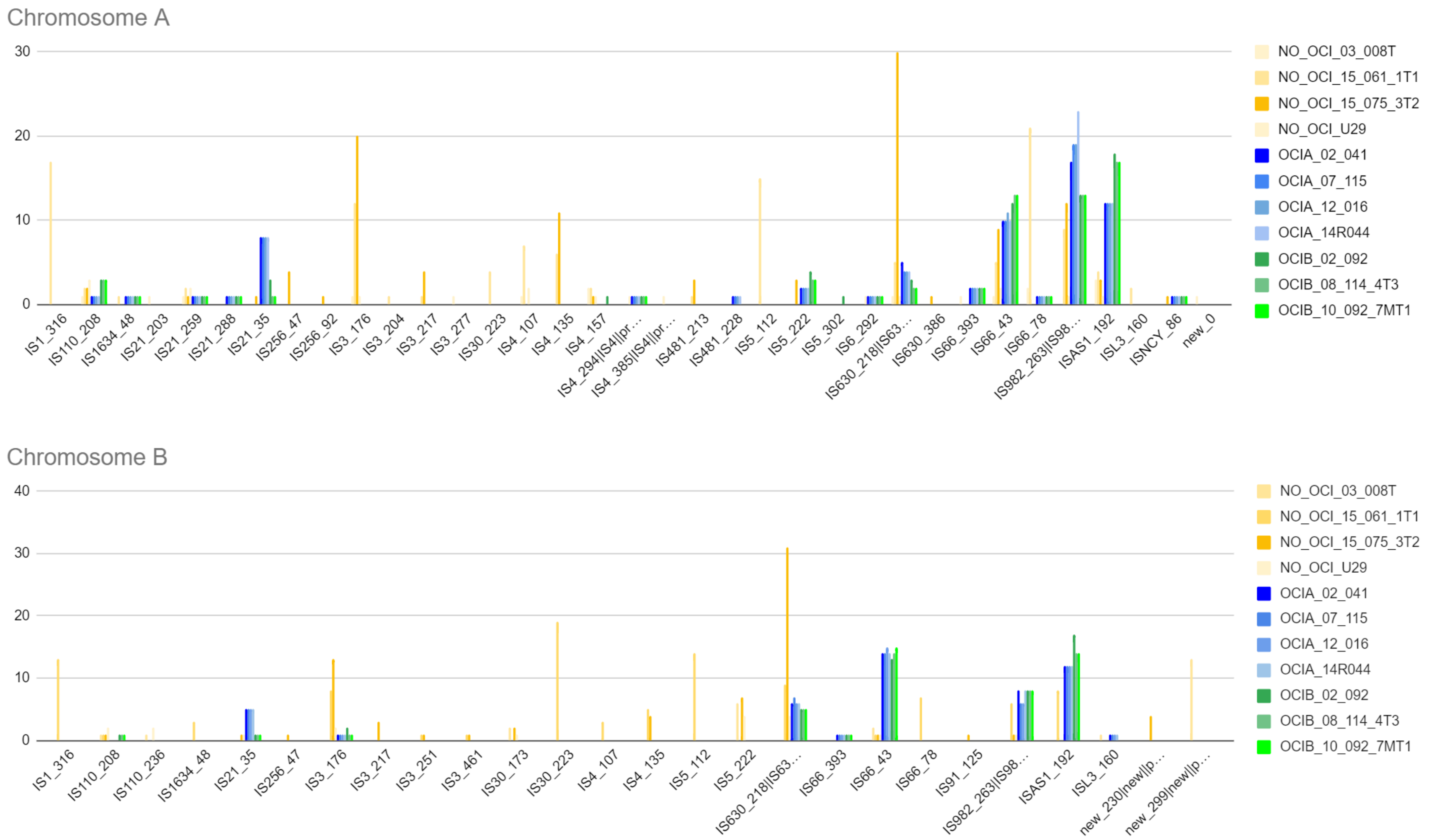


Figure S7: IS number and subgroups on each *Vibrio aestuarianus* chromosome (A and B) for 11 strains other than *Va francensis* (N=4), *Va francensis*, lineage A (N = 4) and B (N=3).

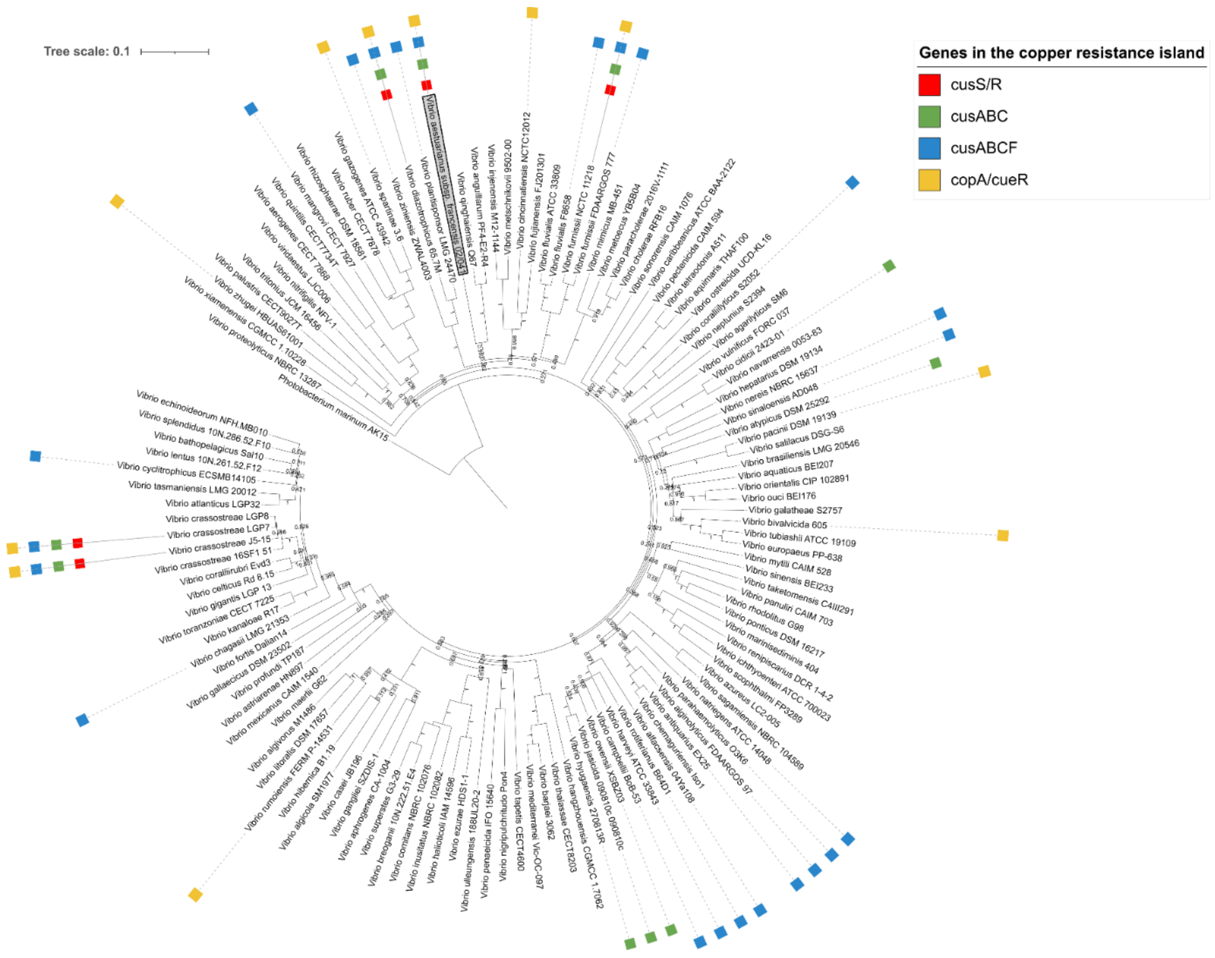


Figure S8: Genes of the copper resistance island occurrence in a same genomic region in 128 vibrio strains. *Va francensis* strain 02_041 is highlighted.
